# Treatment with IgM-enriched intravenous immunoglobulins (IgM-IVIg) enhances clearance of stroke-associated bacterial lung infection

**DOI:** 10.1101/2021.12.16.472965

**Authors:** Laura McCulloch, Alison J. Harris, Alexandra Malbon, Michael J. D. Daniels, Mehwish Younas, John R. Grainger, Stuart M. Allan, Craig J. Smith, Barry W. McColl

## Abstract

Post-stroke infection is a common complication of stroke that is associated with increased mortality and morbidity. We previously found that experimental stroke induces an ablation of multiple sub-populations of B cells and reduced levels of IgM antibody that coincide with the development of spontaneous bacterial pneumonia. Reduced circulating IgM concentrations were also observed in acute stroke patients. The loss of IgM antibody after stroke could be an important determinant of infection susceptibility and highlights this pathway as an important target for intervention.

We treated mice with a low (replacement), dose of IgM-enriched intravenous immunoglobulin (IgM-IVIg) prior to and 24 h after experimental stroke induced by middle cerebral artery occlusion (MCAO) or sham surgery, then recovered mice for 2 d or 5 d. The effect of treatment on lung bacterial burden, lung pathology, brain infarct volume, antibody levels and both lung and systemic cellular and cytokine immune profiles was determined. Treatment with IgM-IVIg enhanced bacterial clearance from the lung after MCAO and improved pathology but did not impact infarct volume. IgM-IVIg treatment induced immunomodulatory effects systemically including rescue of splenic plasma B cell numbers and endogenous mouse IgM and IgA circulating immunoglobulin concentrations that were reduced by MCAO, and treatment also reduced concentrations of pro-inflammatory cytokines in the lung. The effects of MCAO and IgM-IVIg treatment on the immune system were tissue specific as no impact on B cells or mouse immunoglobulins were found within the lung. However, the presence of human immunoglobulins from the IgM-IVIg treatment led to increased total lung immunoglobulin concentration. IgM-IVIg treatment did not increase the number of lung mononuclear phagocytes or directly modulate macrophage phagocytic capacity but enhanced their capability to phagocytose *Staphylococcus aureus* bioparticles *in vitro* by increasing opsonisation.

Low dose IgM-IVIg contributes to increased clearance of spontaneous lung bacteria after MCAO likely via increasing availability of antibody in the lung to enhance phagocytic activity. Immunomodulatory effects of IgM-IVIg treatment, including reduced pro-inflammatory cytokine production, may also contribute to reduced levels of damage in the lung after MCAO. IgM-IVIg shows promise as an antibacterial and immunomodulatory agent to use in the treatment of post-stroke infection.

**GRAPHICAL ABSTRACT:** 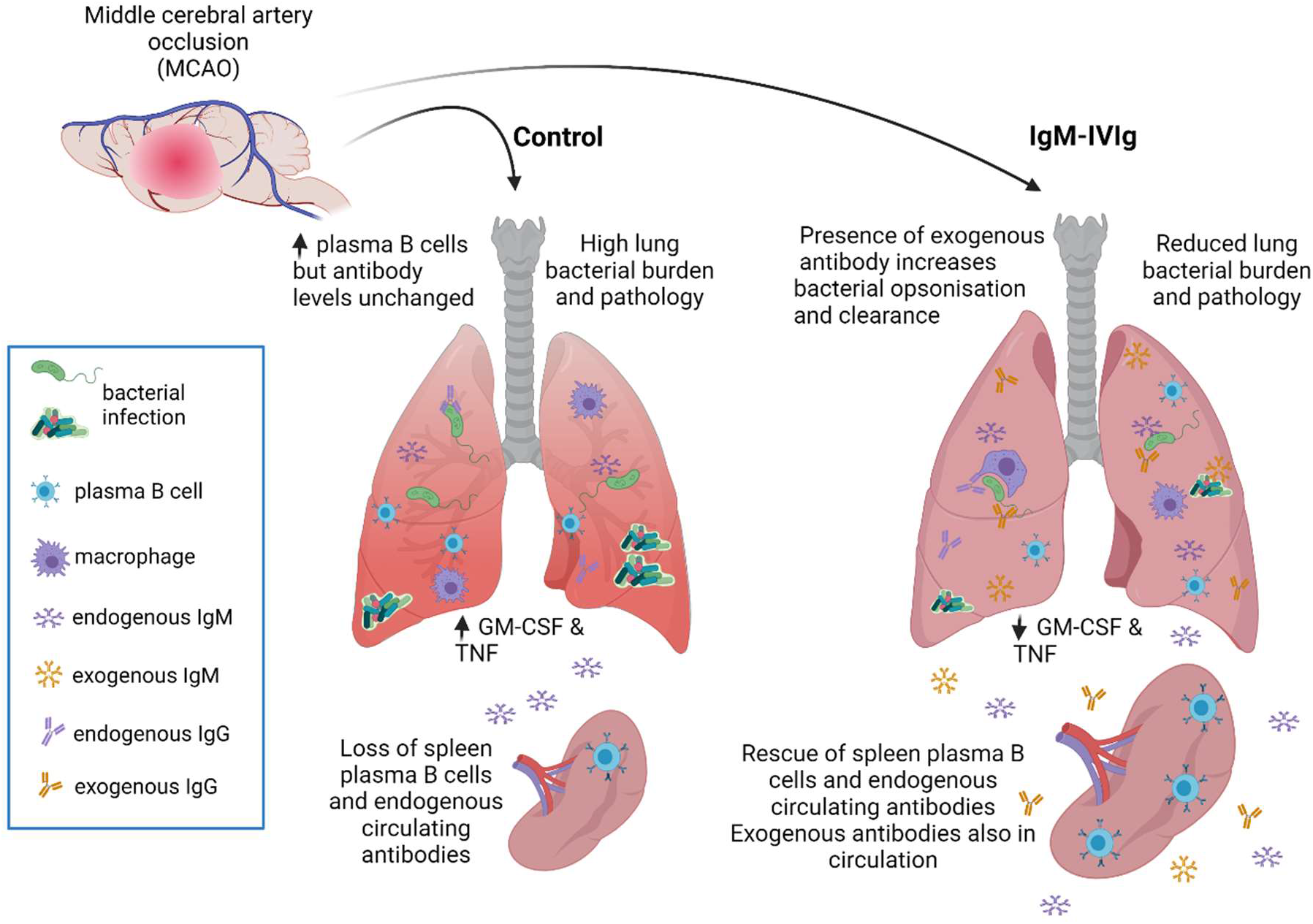

## INTRODUCTION

Improvements in hyper acute stroke care and recent advances made using thrombectomy for removal of large vessel clots have led to an extensive reduction in mortality from ischaemic stroke ^1^. However recovery of patients is often compromised by physical disability, mental health issues and cognitive impairment ^2–4^. Infection is also a serious complication of stroke affecting up to 30 % of patients and infections in both the acute and chronic phase of stroke recovery are associated with increased mortality and disability, and cognitive decline ^5–8^. Indeed, the impact of infection on stroke recovery is so extensive that preventing complications of stroke, such as pneumonia, was listed in the top 10 priorities for stroke research in a recent report from the Stroke Priority Setting Partnership (SPSP)^9^.

Ischaemic stroke induces rapid immune suppression which is thought to contribute to infection risk in patients ^10,11^. Our previous studies identified an early loss of B cells, including marginal zone (MZ) B cells, after experimental stroke with an associated reduction in circulating concentration of IgM antibodies, and the development of spontaneous bacterial pneumonia ^12^. Reduced IgM concentrations were similarly measured in blood samples from patients within one week of ischaemic stroke, compared to controls of similar age and sex distribution ^12,13^. MZ B cell production of IgM provides crucial early defence against systemic infection, in particular infections caused by encapsulated bacteria ^14,15^, which are implicated in pneumonia complicating stroke ^16^. Therefore, reductions in B cells and their production of antibody may be an important determinant of infection susceptibility after stroke.

Intravenous immunoglobulin (IVIg) is a therapeutic agent prepared from pooled serum from thousands of healthy blood donors. Low (replacement) doses of IVIg intended to restore natural circulating antibody concentrations, can be used to treat both primary and secondary immunodeficiencies ^17,18^. At higher doses, IVIg has potent anti-inflammatory properties and has been beneficial in the treatment of autoimmune diseases including Kawasaki disease, Guillain-Barré syndrome and chronic inflammatory demyelinating polyneuropathy (CIDP) ^19,20^. IgM is the most potent antibacterial antibody isotype as its multi-valent structure results in high antigen avidity and LPS-neutralising abilities and potent agglutination and complement fixing properties ^21–23^. IVIg has been used in experimental animal models to treat bacterial pneumonias ^24,25^. Furthermore, IgM-enriched preparations of intravenous immunoglobulin (IgM-IVIg) showed improved efficacy in reducing bacteraemia in animal models of pneumonia and sepsis ^26,27^ and have progressed to clinical trials with improved outcome reported in specific patient sub-groups ^28–30^. Given the antibacterial properties of IgM, and the depleted circulating IgM concentration in patients after acute ischaemic stroke, IgM-IVIg may be a useful therapeutic agent for pneumonia complicating stroke.

Here we show that treatment with IgM-IVIg enhanced the clearance of bacteria and reduced infection-associated pathology in the lung of animals after experimental stroke. Treatment modulated stroke-induced immune alterations, including a rescue of plasma B cells and immunoglobulins systemically and a reduction in lung pro-inflammatory cytokine concentration. The increased availability of human antibodies in the lung provided by IgM-IVIg treatment likely contributes to enhanced phagocytosis of bacteria by increasing opsonisation, as we have demonstrated *in vitro*. This suggests that both direct effect of IgM-IVIg on bacteria and indirect effects modulating the immune environment, contribute to the reduced infectious burden in IgM-IVIg treated mice after MCAO. These studies suggest that IgM-IVIg may be a promising adjunct therapeutic strategy to lessen the impact of pneumonia in patients after stroke.

## MATERIALS AND METHODS

### Animals

Male 8–12-week-old C57BL/6 J mice (Charles River Laboratories, UK) were used in all experiments. Mice were maintained under specific pathogen free (SPF) conditions and a standard 12 h light/dark cycle with unrestricted access to food and water. Mice were housed in individually ventilated cages in groups of up to 6 and were acclimatized for a minimum of 1 week before procedures. All animal experiments were carried out under the authority of a UK Home Office Project Licence in accordance with the ‘Animals (Scientific procedures) Act 1986’ and Directive 2010/63/EU and were approved by the University of Edinburgh’s Animal Welfare and Ethics Review Board. Experimental design, analysis and reporting followed the ARRIVE 2.0 guidelines ^31^. The IMPROVE guidelines were followed to refine procedures and improve animal welfare after MCAO ^32^.

### Experimental stroke model

Middle cerebral artery occlusion (MCAO) surgery was performed under isoflurane anaesthesia induced at 3% and maintained at 1.4% with O_2_ and N_2_O by insertion of a 6-0 nylon monofilament with a 2-mm silicone coated tip (210 μm diameter; Doccol) through the external carotid artery and advanced through the internal carotid artery to occlude the MCA. The filament was withdrawn after 30 min to allow reperfusion, the neck wound sutured and the animals recovered. Topical local anaesthetic (Lidocaine 4%) was applied to the wound. For sham surgery, the filament was advanced to the MCA and immediately retracted. Sham-operated animals remained anaesthetized for 30 min and recovered as above. Core body temperature was maintained at 37±0.5 °C throughout the procedure with a feedback controlled heating blanket (Harvard Apparatus). Animals were recovered for 2 or 5 d then euthanised using a rising concentration of CO_2_ followed by exsanguination.

### Treatment

IgM-IVIg (Pentaglobin; Biotest) was aliquoted neat and given to animals at 250 mg/ kg body weight. HSA (Biotest) was given to animals at 50 mg/ kg. Dilution of control was advised by manufacturer to replicate the use of HSA in clinical trials, where a 5 x lower concentration is used to mimic the foaming, colour and viscosity of IgM-IVIg and therefore assists with protecting blinding when used as a placebo. Treatments were numbered and randomly assigned to animals allowing the surgeon to be blinded to the treatment throughout procedures and welfare assessment during recovery. Treatment was administered intravenously, prior to MCAO or sham surgery and 24 h post-surgery.

### Tissue harvests

Tissue harvests were carried out in a laminar flow cabinet under aseptic conditions. Following euthanasia, the peritoneal cavity was carefully opened and a 23G needle with a 1cm length of sterile polyethylene tubing (Thermo Fisher) over tip was inserted through the diaphragm. Dulbecco’s Phosphate Buffered Saline (800 μl; dPBS; Thermo Fisher) was flushed into the pleural cavity then retrieved using a 1 ml syringe and repeated twice more to obtain a pleural cavity lavage sample. A cardiac blood sample, anti-coagulated with 3.8% w/v tri-sodium citrate, was taken. Half of the sample was stored on ice for bacteriological analysis, the remaining half was spun at 400 x G for 10 min, plasma removed and stored at −80 °C for analysis of soluble mediators. A clean 23G needle with polyethylene tubing was inserted into the trachea and tied in place. dPBS (800 μl) was flushed into the lungs and retrieved using a 1 ml syringe to obtain a bronchial-alveolar lavage (BAL) sample. BAL samples were centrifuged at 800 x G for 10 min and supernatant stored as BAL fluid (BALF). Lungs, heart and thymus were removed from the thoracic cavity as one, with the needle still tied in the trachea. The inferior lobe of the right lung was tied off and removed into a gentleMACS™ C tube (Miltenyi Biotech) containing 1 ml of sterile dPBS for analysis of bacterial load. The middle and superior lobes of the right lung were tied off and removed into a cryovial and snap frozen on dry ice. 800 μl of 4% buffered paraformaldehyde (PFA) was used to inflate the left lung lobe via a needle in the trachea and intact heart, thymus, lung were dropped into a bijou tube of 4 % PFA. After 24 h of immersion fixation, the left lung lobe was dissected from tissue, processed and routinely embedded in paraffin blocks for histology. Next the spleen was dissected, weighed and halved. Half was snap frozen in a cryovial over dry ice for immunostaining whilst the other half was taken into a bijou tube and stored on ice for flow cytometry. Finally, the brain was removed and frozen in cold isopentane over dry ice and stored in a bijou tube at −80 °C.

Flow cytometry of the lungs at 2 d post sham or MCAO surgery was carried out in a separate cohort of animals. Animals were euthanised in a rising concentration of CO_2_ followed by exsanguination. Pleural cavity lavage was taken as previously described. BAL was also taken as above, however the lungs were flushed three times with 800 μl dPBS to maximise cellular harvest. The entire lung was dissected into 1 ml cold RPMI 1640 medium (RPMI; Thermo Fisher). Samples were stored on ice until processed for flow cytometry.

### Exclusion criteria

Post-hoc exclusions of tissues for analysis were applied as follows. Animals that did not have infarcts affecting the cortex and striatum after MCAO surgery were completely excluded from the study as an unsuccessful MCAO (n=5). For lung pathology, lungs that were not fully inflated before fixation and showed crush artefact were excluded from analysis (2 d Sham IgM-IVIg n=1; MCAO HSA n=1; 5 d HSA n=1; IgM-IVIg n=3). Plasma samples from failed cardiac punctures were excluded from multiplex ELISA analysis (2 d Sham IgM-IVIg n=2; MCAO HSA n=1). BALF samples which had low/ undetectable levels of protein were excluded from multiplex ELISA analysis (2 d Sham HSA n= 2; IgM-IVIg n=3; MCAO HSA n=2; MCAO IgM-IVIg n=1). Samples were excluded from analysis of spleen flow cytometry where staining of one or more antibodies in the cocktail failed (2 d Sham HSA n= 1; IgM-IVIg n=1; MCAO HSA n=2; MCAO IgM-IVIg n=1). Lung samples with cell harvest < 1 x 10 7 total cells or where staining of one or more antibodies in the cocktail failed were excluded from flow cytometry analysis failed (2 d Sham HSA n= 2; IgM-IVIg n=1; MCAO HSA n=2; MCAO IgM-IVIg n=1).

### Infarct quantification

Coronal cryosections (20 μm) were taken at 400 μm intervals from fresh frozen brains and stained with Harris Haematoxylin stain-acidified (CellPath) and Eosin Y (CellPath) (H&E). Briefly, sections were hydrated by soaking for 2 min each in 100% > 95% > 80% > 70% ethanol, then distilled water. Slides were stained with haematoxylin for 5 min, washed in distilled water, immersed in Scott’s tap water (CellPath) for 30 s followed by a 30 s wash in distilled water. Sections were then stained with eosin for 3 min and washed with distilled water. Sections were dehydrated by soaking for 2 min in 70% > 80% > 95% > 100% ethanol, fixed for 5 min in xylene, then coverslips were applied using DPX mountant (Merck). Sections were scanned on an Axio Scan.Z1 (Zeiss) and visualized using Zeiss Zen 3.2 blue edition software. Sections identified by neuroanatomical landmarks at 8 specific coronal levels for quantification as described in ^33^. The infarcted area for each region (cortex, striatum/pallidum, hippocampus, thalamus and other) at each of the 8 coronal levels was quantified using Fiji (U. S. National Institutes of Health, USA https://imagej.nih.gov/ij/) and summed to give a total for each section. These values were plotted against distance from rostral pole and area under the curve calculations performed in Prism (GraphPad Prism v9) to calculate infarct volume.

### Quantifying lung bacterial load

The inferior lobe of the right lung was homogenised in 1 ml sterile dPBS using a gentleMACS (Miltenyi Biotech). In a laminar flow hood, lung homogenates and blood, collected as described above, were serially diluted from neat to 10^−6^ in sterile dPBS and dilutions plated onto Columbia agar with horse blood for a general bacteria screen or MaConkey’s agar to assess gut-derived bacteria. Plates were incubated at 37 °C for 48 h and bacterial colonies counted.

### Lung pathology

Sections (4 μm) were cut from inflated left lungs embedded in paraffin wax blocks. Sections were dewaxed in xylene and then stained with H&E as above. Pathology was assessed by a board-certified veterinary pathologist, who was blinded to treatment groups, using scoring criteria described in **Supplementary Figure 2**.

### Multiplex ELISA

Human and mouse immunoglobulins and mouse cytokines were measured in plasma samples and BALF using MILLIPLEX® multiplex assays. Coded samples were randomised across plates for analysis to ensure the researcher was blinded to treatment group. The MILLIPLEX®MAP Human Isotyping Magnetic Bead Panel-Isotyping Multiplex Assay (HGAMMAG-301K-06, Merck) was used to measure IgG1, IgG2, IgG3, IgG4, IgA and IgM. Plasma samples were diluted 1/ 16,000 before adding to plate. Total protein concentration of BALF was measured using a Pierce™ BCA Protein Assay Kit (Thermo Fisher). BALF volume equivalent to 25 μg of protein was added to each well to normalise across groups accounting for any effects of imbalances in protein concentration due to treatments. MILLIPLEX®MAP Mouse Isotyping Magnetic Bead panel (MGAMMAG-300K, Merck) was used to measure IgG1, IgG2a, IgG2b, IgG3, IgA, IgM. Many samples had concentrations of IgG2a below the detection range of the standard curve and so results for this analyte are not reported. Plasma samples were added to plate at a 1/25,000 dilution and BALF samples were added to plate as previous. MILLIPLEX MAP Mouse Cytokine/Chemokine Magnetic Bead Panel (MCYTOMAG-70K, Merck) was used to measure GM-CSF, IFN γ, IL-1α, IL-1β, IL-2, IL-4, IL-5, IL-6, IL-12(p40), IL-33 and TNF α. Many samples had concentrations of IL-1β below the detection range of the standard curve and so results for this analyte were not reported. Plasma samples were added to plate at a 1/10 dilution and BALF samples were added to plate as previous. In all assays, samples were assayed as single replicates and all samples, standards and quality controls were prepared in accordance with the manufacturer’s instructions. Samples were incubated with beads on a plate for 1 h (isotyping assay) at room temperature or overnight (cytokine assay) at 4° C and washes carried out using a magnetic plate washer. Plates were analysed using a Magpix™ Luminex® machine and Luminex xPonent® software version 4.2, with a sample volume of 50 μl per well and a minimum of 50 events counted per sample.

### Flow cytometry

Spleens were disrupted with a syringe plunger and passed through a 100 μM cell sieve. Red blood cells were lysed with red blood cell lysing buffer Hybri-MaxTM (Sigma) before counting live cells with trypan blue (Sigma) exclusion. Splenocytes were plated at 2 million cells/well and incubated with Zombie UV (Biolegend) for dead cell discrimination, then Fc blocked with normal mouse serum (Thermo Fisher) and TruStain FcX αmouse CD16/32 (Biolegend) before staining with antibody cocktails. B cell subsets and other lymphoid and myleloid populations were identified using combinations of the following anti-mouse antibodies: CD3 PerCP-Cy5.5, CD3 BV785, CD11b BV711, CD11c AF488, CD20 PerCP-Cy5.5, CD24 PerCP-Cy5.5, CD43 PE-Cy7, CD45 BV510, CD64 PE, I-A/I-E (MHCII) BV785, IgD BV711, IgM BV421, Ly6C APC-Cy7, Ly6G PE-Cy7, NKp46 PE/Dazzle594 (all Biolegend); CD19 BUV395, CD21/35 PE-CF594, CD23 BB515, SiglecF AF647 (all Becton Dickinson); CD5 APC, CD93 APC, CD138 PE (all Miltenyi Biotec).

Lungs were removed from RPMI, chopped with scissors and 2 ml of enzyme cocktail [0.8 mg/ml Collagenase V (Sigma), 0.625 mg/ml Collagenase D (Roche), 1 mg/ml Dispase (Gibco), 30 μg/ml DNase (Roche) in RPMI] added before 45 min incubation at 37°C on a shaking incubator. Digested lung homogenate was passed through a 100 μm cell strainer. Red blood cells were lysed with red blood cell lysing buffer Hybri-MaxTM (Sigma) before counting live cells with trypan blue (Sigma) exclusion. Lung cells were plated at 2 million cells/well and incubated with Zombie UV (Biolegend) for dead cell discrimination, then Fc receptors blocked with normal mouse serum (Thermo Fisher) and TruStain FcX αmouse CD16/32 (Biolegend) before staining with antibody cocktails. B cell subsets and other lymphoid and myeloid populations were identified using combinations of the following anti-mouse antibodies: CD3 BV785 or PerCP-Cy5.5, CD11b BV711, CD20 BV421, CD43 PE-Cy7, CD45 BV510, CD64 PE, Ly6C APC-Cy7, Ly6G PE-Cy7, MHC II (I-A/ I-E) BV785 or APC-Cy7, NKp46 PE/ Dazzle 594 (all Biolegend); CD5 APC (Miltenyi Biotec), CD11c BUV395 and Siglec F AF647 (all Becton Dickinson).

Cells were fixed using a formaldehyde-based fixation buffer (Biolegend) and analysed within 48 h on a 5-laser LSR Fortesssa (Becton Dickinson). Data analysis was performed on Flowjo (Treestar), with FSCA/FSCH doublet exclusion and dead cell exclusion. Spleens were weighed then divided into two in order to perform both flow cytometry and IHC. The weight of each half as a proportion of the total weight was used to convert the cell count for flow cytometry into an estimated total for the whole spleen as follows:

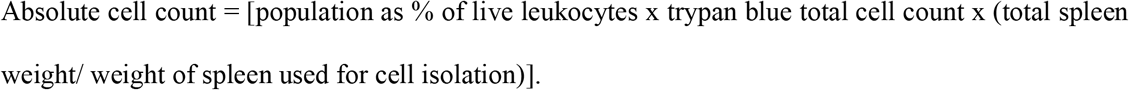

### Spleen immunohistochemistry

Acetone fixed 6 μm spleen sections were blocked with normal goat serum (Sigma). For detection of germinal centres, sections were labelled with rat anti-mouse B220 (Biolegend RA3-6B2) followed by goat anti-rat AF594 secondary antibody (Thermo Fisher). Sections were washed and subsequently incubated with AF647-conjugated lectin peanut agglutinin (PNA; Thermo Fisher) in the presence of sodium azide and bovine serum albumin overnight at 4°C.

### *In vitro* phagocytosis assay

Bone marrow single cell suspensions were made by flushing cells from the femurs and tibias of C57Bl/6J mice into sterile dPBS. Cells were resuspended in RPMI containing 10% foetal bovine serum (Thermo Fisher) and 1% Penicillin-Streptomycin (Thermo Fisher) and 25% conditioned media from L929 cells and incubated at 37°C with 5% CO_2_. Cultures were supplemented with fresh medium at 24 h. After 7 d, differentiated bone marrow-derived macrophages (BMDM) were transferred to a 96 well plate at 5 ×10^5^ cells/ ml and incubated for 24 h prior to use in assay. Cells were either untreated or given treatment 0.25 mg/ml or 2 mg/ ml IgM-IVIg, control 0.05 mg/ml or 0.4 mg/ml HSA and returned to incubator for 4 h. After incubation with treatment, cells were either “washed” in PBS with fresh RPMI replaced or left “unwashed” and proceeded directly to next step of assay. Plate was inserted into an IncuCyte® S3 Live-Cell Analysis System (Sartorius) for a single phase-contrast read in order to determine cell count. 10 μM Cytochalasin D was added to negative control wells for 10 min, then pHrodo® Red *Staph. aureus* BioParticles® (Thermo Fisher) were diluted 1:4 in RPMI and 10 μl added to each well of cells. Plates were then returned to the IncuCyte® and imaged every 5 min for 2.5 h on both phase-contrast and ‘Red’ channels (400 ms exposure time). Cell-by-cell analysis was performed on Incucyte® analysis software, using a mask to automatically identify (1) all BMDM, using phase contrast and (2) “red objects” i.e. BMDM that have phagocytosed pHrodo® Red bioparticles. Thresholds for size and fluorescence were adjusted during a pilot experiment, to eliminate non-specific signal while maximising detection of phagocytosing cells, with visual inspection of multiple FOVs to check sensitivity and accuracy. Integrated density of red fluorescent signal was measured at each time point and was normalised to the total number of macrophages present in the well at the beginning of assay.

### Experimental design and statistical analysis

Sample sizes were estimated from previous data on reduced IgM concentration and reduced splenic MZ B cells after experimental stroke ^12^ using power analysis carried out on InVivoStat software (http://invivostat.co.uk/) to be sensitive to a 35% effect of experimental stroke at 80% statistical power at a 5 % significance level. Animals were ear notched for identification and randomised to experimental groups (sham or MCAO surgery and treatment) using a computer-based random number generator (https://www.randomizer.org/) and treatments were administered in a blinded manner and allocation of treatment was concealed throughout study. Animals were given an experimental identifier and all samples were analysed using this coded identifier. Data was unblinded for analysis after experimental work was complete.

For normally distributed data, differences were tested using unpaired Student’s t-test or 2-way analysis variance (ANOVA) with Tukey multiple comparison test. Results were displayed as mean ± standard deviation. When data were non-normally distributed, log transformation was used to normalise data sets and data were then analysed as previous. Non-parametric tests, such as Mann-Whitney test, were used to analyse data produced from scoring systems and results were displayed as violin plots. Data were analysed using GraphPad Prism. In all experiments, values of P≤0.05 were accepted as statistically significant.

## RESULTS

### Low dose IgM-IVIg enhances clearance of spontaneous bacterial infection after experimental stroke

To determine if replacing immunoglobulins depleted after stroke is an effective therapeutic strategy to improve bacterial pneumonia, animals were given a low dose of IgM-IVIg (250 mg/kg) prior to experimental stroke induced by transient MCAO or sham surgery and at 24 h post-surgery. At 2 d post-MCAO there was a trend for mice treated with IgM-IVIg to have reduced general (blood agar; **Figure1A**,) and gut-derived (MacConkey agar; **Figure 1B**) lung bacterial burdens in comparison to HSA treated mice. By 5 d post-MCAO, both general (**Figure 1C**) and gut-derived (**Figure 1D**) lung bacterial burdens were significantly reduced in mice treated with IgM-IVIg. Spontaneous bacterial pneumonia after stroke is known to be associated with stroke severity ^34^. Treatment with IgM-IVIg did not alter brain infarct volume size at 2 d or 5 d post-MCAO (**Supplementary Figure 1A-D**) demonstrating that reduced bacterial counts were a direct effect of the treatment and not due to alterations in primary stroke pathology.

**Figure 1.**
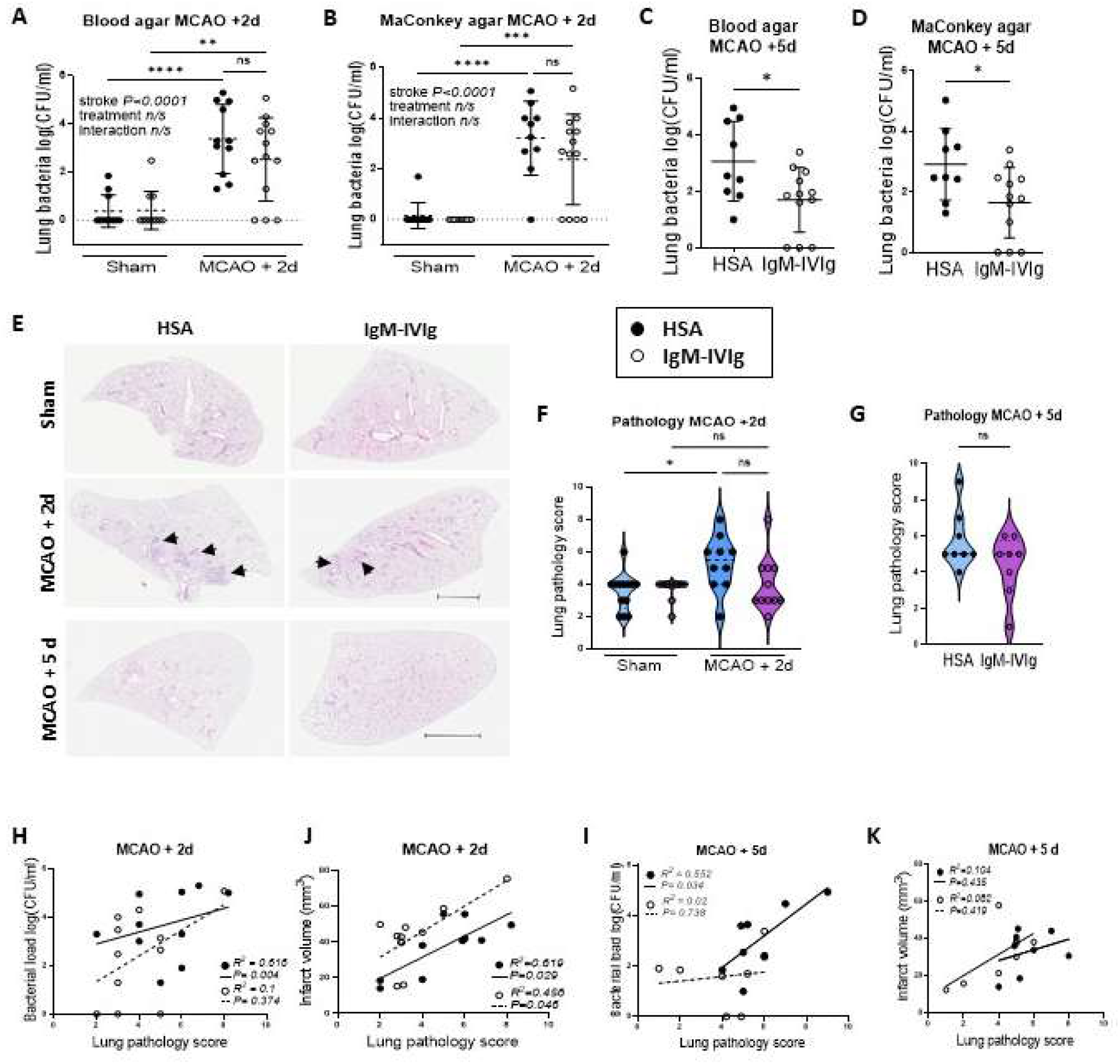
Low dose IgM-IVIg enhances clearance of spontaneous bacterial infection after experimental stroke. Bacterial load (log[CFU/ml]) in the inferior lobe of the right lung in IgM-IVIg (○) or human serum albumin (HSA; ●) treated mice plated on blood agar at (**A**) 2d or (**C**) 5d after MCAO or sham surgery and plated on MacConkey agar at (**B**) 2 d or (**D**) 5 d after MCAO or Sham surgery (Sham HSA n=12; Sham IgM-IVIg n=11; MCAO HSA n=11; MCAO IgM-IVIg n=13). (**E**) Representative haematoxylin and eosin stained sections of the left lung 2 d after sham or MCAO surgery in IgM-IVIg or HSA treated mice. Arrows denote inflammatory cluster. Pathology score of left lung at (**F**) 2 d or (**G**) 5 d after sham or MCAO surgery in IgM-IVIg (○) or HSA (●) treated mice from scoring criteria detailed in Supplementary Fig 2 (Sham HSA n=12; Sham IgM-IVIg n=9; MCAO HSA n=10; MCAO IgM-IVIg n=10). The association between lung pathology score and lung bacterial load (log [CFU/ml]) at (**H**) 2 d and (**I**) 5 d post MCAO in IgM-IVIg (○) or HSA (●) treated mice. The association between lung pathology and stroke infarct volume at (**J**) 2 d and (**K**) 5 d post MCAO in HSA (●) and IgM-IVIg (○) treated mice. Data show data points with mean ± S.D; * P<0.05; (A, C) two-way ANOVA with Tukey’s multiple comparison test; (B, D) unpaired t-test. (F, G) Violin plots with Mann-Whitney test. (H-K) Simple linear regression with Pearson’s correlation.

Lung pathology was assessed using a scoring system that accounted for perivascular edema, inflammation in the perivascular and peribronchiolar regions, macrophages in the alveolar spaces and macrophages/ hypercellularity in the interstitium (**Supplementary Figure 2**). At 2 d post-surgery, a low level of lung pathology was seen in sham-operated animals, regardless of treatment, which may be due to the time under inhalational anaesthesia during surgery. At 2 d post-MCAO, lung pathology scores were significantly increased in HSA treated animals, whereas in IgM-IVIg treated animals there was no difference to sham-operated controls (**Figure 1E, F**). A trend for reduced lung pathology scores in IgM-IVIg treated animals was also observed at 5 d after MCAO (**Figure 1E, G**).

Associations between lung pathology and bacterial burden, or stroke severity, were explored using these data. There was a modest association between lung pathology and bacterial burden at 2 d and 5 d post MCAO (**Figure 1H, I**) and between lung pathology and infarct volume at 2 d post MCAO in HSA treated animals. The association between lung pathology and bacterial burden was not observed in IgM-IVIg treated animals but there was an association between lung pathology and infarct volume at 2 d post MCAO. Consistent with previous data showing that IgM-IVIg did not alter brain infarct volume, this indicates that IgM-IVIg treatment modulates the association between bacterial burden and lung pathology observed in HSA treated controls. Furthermore, as bacterial burden and lung pathology were altered to different degrees in IgM-IVIg treated animals, some lung pathology may be driven by the stroke itself, in addition to that caused by infection.

Although high-dose IVIg treatment is typically associated with inhibitory immunoreceptor signalling, IVIg have a range of known immunomodulatory properties, including the potential to also trigger activating Fc receptor pathways. There was a possibility that this treatment could have detrimental effects on recovery from stroke (e.g. if harmful hyper-immune responses were triggered). No differences in general welfare score (**Supplementary Figure E, F**) or neurological deficit (Bederson Score; **Supplementary Figure 1G, H**) were observed at either time point. Treatment with IgM-IVIg did result in higher body weight loss 2 d after both sham and stroke surgery (**Supplementary Figure 1I**) however this normalised by 5 d post-stroke (**Supplementary Figure 1J**). There was no difference in mortality between treatment groups at 2 d post-stroke, however reduced mortality was seen in IgM-IVIg treated mice at 5 d (**Supplementary Figure 1K**).

These data demonstrate that low dose IgM-IVIg enhances the clearance of spontaneous bacterial infection after experimental stroke. IgM-IVIg treatment led to reduced lung pathology in comparison to HSA treated animals with equivalent bacterial burden or stroke severity. Furthermore, although an early increase in weight loss is seen in IgM-IVIg treated animals, there were no adverse effects of IgM-IVIg treatment on primary stroke pathology, welfare or mortality.

### Distribution of human immunoglobulins in IgM-IVIg treated animals

We next wanted to determine that IgM-IVIg treatment was present both systemically, and within the lung, after intravenous administration. As IgM-IVIg is pooled from human donors, multiplex ELISA to detect human immunoglobulins can be used to distinguish the presence of IgM-IVIg from endogenously produced mouse immunoglobulin. Human immunoglobulin subsets IgM, IgG1, IgG2, IgG3, IgG4 and IgA were significantly increased in the plasma of animals receiving IgM-IVIg treatment at 2 d post MCAO or sham surgery (**Figure 2A-F**). Low levels of some immunoglobulin isotypes were detected in HSA treated animals, indicating a mild cross reactivity of assay with endogenous mouse immunoglobulin, however the presence of human immunoglobulins were clearly detected above this background in IgM-IVIg treated animals (**Figure 2A-F**). Human immunoglobulins were also detected in BALF, although at relatively lower concentrations than detected in plasma (**Figure 2G-L**). The proportion of IgM, IgA and total IgG detected in plasma and BALF was compared to the original IgM-IVIg treatment. There was proportionally less IgM in the BALF than was detected in the plasma or in the original IgM-IVIg treatment (**Figure 2M**). These data demonstrate that IgM-IVIg-derived human immunoglobulins can be detected both systemically, and within the lung, after intravenous administration. However there may be relatively less access of human IgM to the lung compartment than other immunoglobulin isotypes.

**Figure 2.**
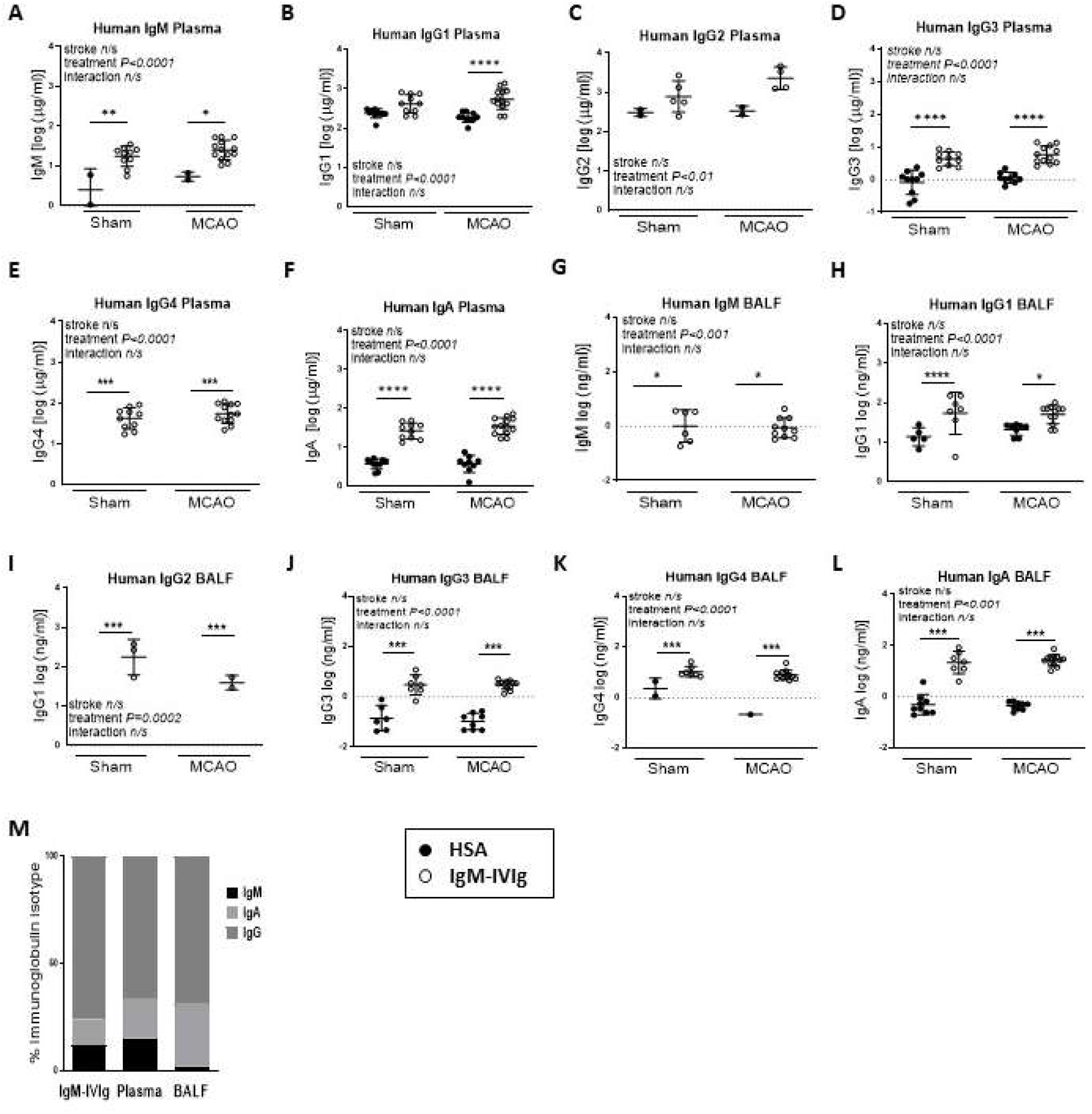
Distribution of human immunoglobulins in IgM-IVIg treated animals. Presence of IgM-IVIg in the circulation was detected by measuring concentration of human immunoglobulin isotypes (**A**) IgM, (**B**) IgG1, (**C**) IgG2, (**D**) IgG3, (**E**) IgG4 and (**F**) IgA in plasma before treatment (baseline) with human serum albumin (HSA; ●) or IgM-IVIg (○) and after 2 d recovery from sham or MCAO surgery (Sham HSA n=11; Sham IgM-IV-Ig D2 n=10; MCAO HSA D2 n=10; MCAO IgM-IVIg D2 n=13). Presence of IgM-IVIg in the lung was detected by measuring concentration of human (**G**) IgM, (**H**) IgG1, (**I**) IgG2, (**J**) IgG3, (**K**) IgG4 and (**L**) IgA in bronchial alveolar lavage fluid (BALF) in animals treated with HSA (●) or IgM-IVIg (○) after 2 d recovery from sham or MCAO surgery (Sham HSA n=9; Sham IgM-IVIg n=8; MCAO HSA n=9; MCAO IgM-IVIg n=12). (**M**) The proportion of human IgM, IgG and IgA reported to be present in IgM-IVIg in comparison to proportion detected in plasma and BALF. Data show data points with mean ± S.D; * P<0.05; ** P<0.01; *** P<0.001; (A-L) two way ANOVA with Tukey’s multiple comparison test.

### Low dose IgM-IVIg rescues splenic plasma B cells and endogenous antibody production after experimental stroke

Experimental stroke is known to result in lymphopenia, and in particular a loss of B cells and circulating antibodies, which is thought to contribute to the onset of spontaneous bacterial pneumonias ^12^. Furthermore, IgM signalling via the Fcμ receptor can have autocrine effects on B cell development and activation ^35^. We therefore characterised the effects of experimental stroke on B cell subpopulations and endogenous mouse immunoglobulins and determined if these were altered by treatment with IgM-IVIg. Flow cytometry was used to detect splenic B cells (**Figure 3A**), 2 d after experimental MCAO or sham surgery and treatment with HSA or IgM-IVIg. A reduction in spleen weight was observed 2 d after MCAO as previously reported ^12^ (**Supplementary Figure 3A**). Treatment with IgM-IVIg did not alter spleen weight 2 d or 5 d after MCAO suggesting no generalised effect of treatment on systemic immune cellularity (**Supplementary Fig 3A, B**). In confirmation, MCAO-induced reduction in total spleen leukocytes was also unaltered by treatment with IgM-IVIg (**Figure 3B**). Furthermore, total CD19^+^ splenic B cells were reduced by MCAO, but unaffected by treatment with IgM-IVIg (**Figure 3C**).

**Figure 3.**
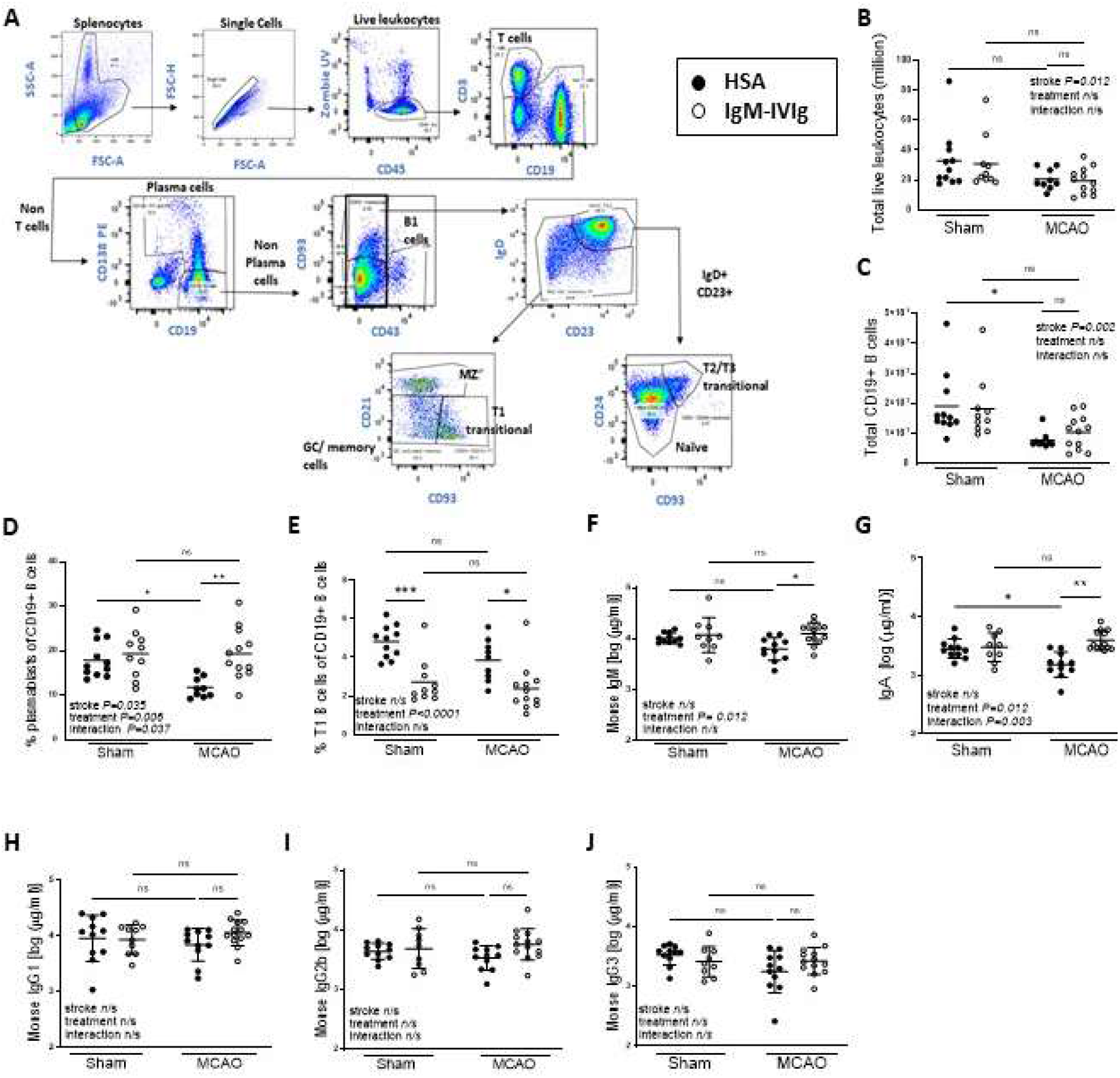
Low dose IgM-IVIg rescues splenic plasma B cells and endogenous antibody production after experimental stroke. (**A**) Gating strategy used to define spleen B cell subsets by flow cytometry. (**B**) Total live leukocytes and (**C**) total CD19^+^ B cells measured by flow cytometry of spleens from mice treated with human serum albumin (HSA; ●) or IgM-IVIg (○) and after 2 d recovery from sham or MCAO surgery. (**D**) The proportion of CD138^+^ plasma cells and plasmablasts and (**E**) CD138^−^CD43^−^CD21^−^CD23^−^CD93^+^ T1 Transitional B cells within the CD19^+^ total B cell population mice treated with human serum albumin (HSA; ●) or IgM-IVIg (○) and after 2 d recovery from sham or MCAO surgery (Sham HSA n=11; Sham IgM-IVIg n=10; MCAO HSA n=9; MCAO IgM-IVIg n=12). Circulating concentration of endogenously produced mouse immunoglobulins (**F**) IgM, (**G**) IgA, (**H**) IgG1, (**I**) IgG2a, (**J**) IgG2b and (**K**) IgG3 in plasma in animals treated with HSA (●) or IgM-IVIg (○) after 2 d recovery from sham or MCAO surgery (Sham HSA n=11; Sham IgM-IVIg n=10; MCAO HSA n=10; MCAO IgM-IVIg n=13). Data show data points with mean ± S.D; * P<0.05; ** P<0.01; *** P<0.001, (B-K) 2 way ANOVA with Tukey’s multiple comparison test.

We next determined if IgM-IVIg affected B cell sub-populations within the CD19^+^ splenic B cell pool. Experimental stroke resulted in a reduction in the proportion of antibody secreting CD138^+^ plasma cells and plasmablasts. Treatment with IgM-IVIg rescued plasma B cell abundance after MCAO (**Figure 3D**). In contrast, the proportion of T1 transitional B cells was unaffected by stroke, but reduced in animals treated with IgM-IVIg (**Figure 3E**). T2/T3 transitional B cells, naïve B cells, pooled GC and memory B cells, marginal zone B cells and innate-like B1 B cells were unaffected by treatment with IgM-IVIg (**Supplementary Figure 3C-G**). There was a trend for more mature germinal centres (GC) to be present in spleens of IgM-IVIg treated animals 2 d after either MCAO or sham, surgery (**Supplementary Figure 3H, I**).

Multiplex ELISA was used to measure endogenous mouse immunoglobulins in plasma. Treatment with IgM-IVIg restored circulating concentration of mouse IgM (**Figure 3F**) and IgA (**Figure 3G**) in animals 2 d after MCAO, whereas concentrations of mouse IgG1, 2b and 3 (**Figure 3H, I, J**) were unaffected by treatment.

In summary, treatment with IgM-IVIg in experimental stroke prevented the MCAO-induced reduction in plasma B cells abundances and circulating IgM and IgA antibody concentrations, reductions in transitional B cells, and had no effect on other antibody secreting cellular subsets (B1 and MZ B cells). Taken together, these data suggest treatment with IgM-IVIG may drive T1 B cells into the GC response producing increased output of plasma B cells and a rescue of endogenous mouse IgM and IgA circulating antibody concentration.

### Low dose IgM-IVIg does not modulate lung B cells or endogenous immunoglobulins in the lung after experimental stroke

As experimental stroke results in spontaneous bacterial pneumonia, we next investigated if similar effects of MCAO and IgM-IVIg treatment occurred in antibody secreting B cell populations and immunoglobulin concentrations within the lung. Flow cytometry was used to detect antibody-secreting B cell populations in the lung (**Figure 4A**). MCAO induced an increase in the number of leukocytes in the lungs of mice at 2 d and this was not altered by treatment with IgM-IVIg (**Figure 4B**). In contrast to splenic B cell numbers (**Figure 3C**), total CD20^+^ B cells were not reduced by MCAO but were reduced by IgM-IVIg treatment in both sham and experimental stroke animals (**Figure 4C**). Furthermore, MHC II^−^ mature plasma cells in the lung increased after MCAO, most likely in response to bacterial infection (**Figure 4D**). A trend for lower levels of plasma cells was seen in the lungs of IgM-IVIg treated mice after MCAO, but not sham, surgery (**Figure 4D**). As there was no effect in sham animals, this may reflect the lower levels of bacterial infection in IgM-IVIg treated animals after MCAO, rather than a direct effect of IgM-IVIg on plasma cell numbers. Innate-like B1 cells can rapidly produce antibody in response to infection and are an additional and important source of antibody in the lung ^36^. B1 B cells were unaltered by either MCAO or IgM-IVIg (**Figure 4E**).

**Figure 4.**
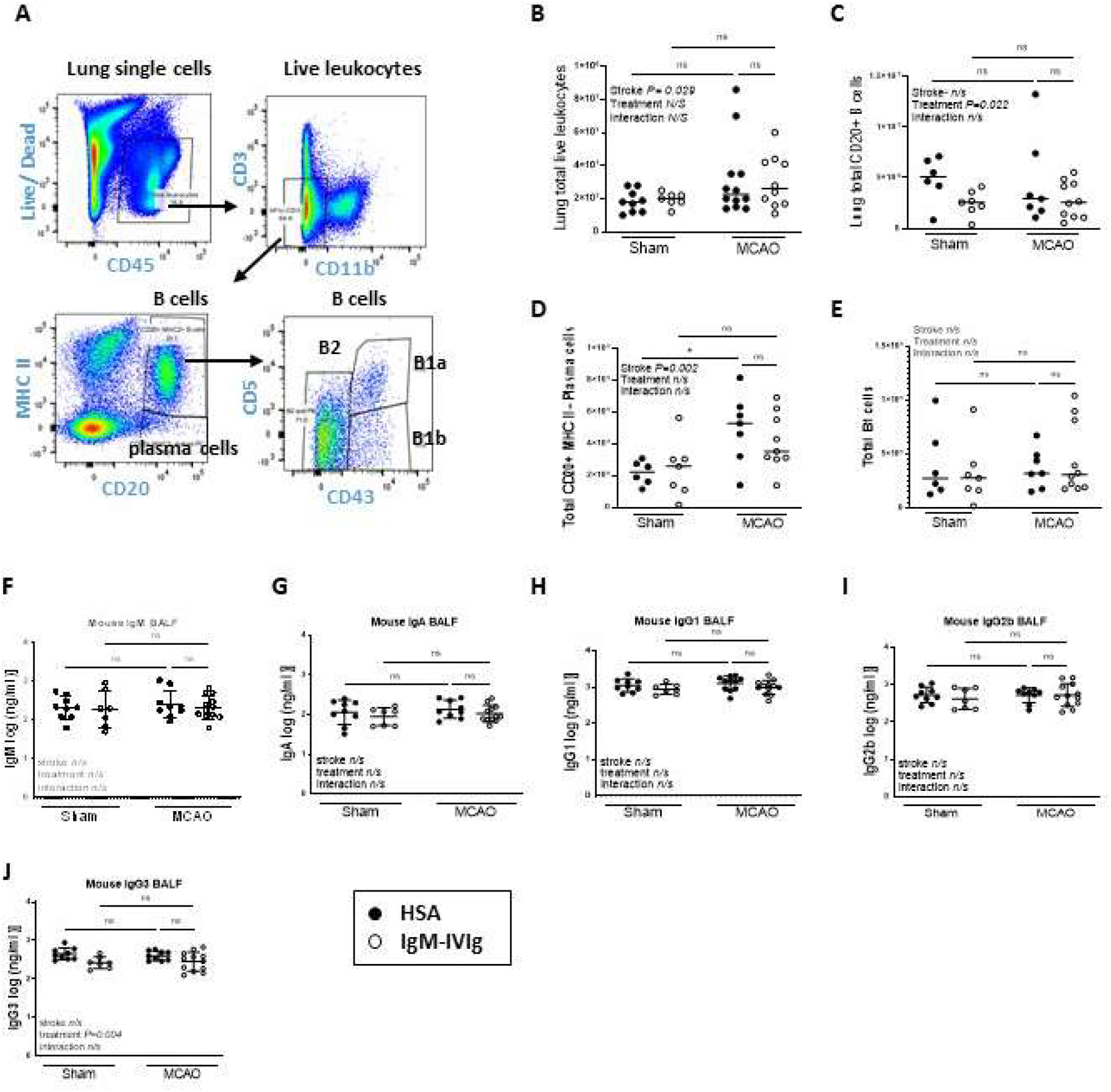
Low dose IgM-IVIg does not modulate lung B cells or immunoglobulins after experimental stroke. (**A**) Gating strategy used to define lung B cell subsets by flow cytometry. (**B**) Total live lung leukocytes, (**C**) CD20^+^ B cells (**D**) CD20^+^MHC II^−^ plasma B cells and (**E**) CD20+MHC II+ CD43+CD5+/-B1 B cells measured by flow cytometry of lung single cell suspensions from mice treated with human serum albumin (HSA; ●) or IgM-IVIg (○) and after 2 d recovery from sham or MCAO surgery (Sham HSA n=6; Sham IgM-IVIg n=7; MCAO HSA n=7; MCAO IgM-IVIg n=10). Concentration of endogenously produced mouse immunoglobulins (**F**) IgM, (**G**) IgA, (**H**) IgG1, (**I**) IgG2b and (**J**) IgG3 and in bronchial alveolar lavage fluid (BALF) in animals treated with HSA (●) or IgM-IVIg (○) after 2 d recovery from sham or MCAO surgery (Sham HSA n=9; Sham IgM-IVIg n=8; MCAO HSA n=9; MCAO IgM-IVIg n=12). Data show data points with mean ± S.D; * P<0.05; (B-J) two way ANOVA with Tukey’s multiple comparison test.

We next examined the effects of MCAO and IgM-IVIg treatment on endogenous mouse immunoglobulin concentrations in the lung. Despite a MCAO-induced increase in plasma cells within the lung, there was no proportional increase in mouse IgM, IgA, IgG1, IgG2b or IgG3 in the BALF (**Figure 4F, G, H, I, J**). There was also no additional effect of treatment with IgM-IVIg, with the exception of reduced concentration of IgG3 in sham-operated animals only (**Figure 4J**). This contrasts with circulating immunoglobulin concentrations where reduced levels of IgM and IgA after MCAO are restored by treatment with IgM-IVIg (**Figure 3F, G**).

These data demonstrate the differential effects of MCAO and IgM-IVIg on endogenous mouse cellular and immunoglobulin profiles within the lung versus systemically, as measured in the spleen. Whereas systemically, MCAO reduces B cell numbers, antibody-secreting plasma cell numbers and circulating IgM and IgA, there was no reduction in B cells or antibody concentrations observed in the lung. There was an increase in plasma cells in the lung after MCAO but this was not accompanied by an increase in immunoglobulin concentrations which may indicate some MCAO-induced functional deficits. Furthermore, treatment with IgM-IVIg prevented a loss of splenic plasma cell numbers and IgM and IgA concentration, but has no effect on lung plasma cells or immunoglobulins. Despite the lack of effect of IgM-IVIg treatment on mouse immunoglobulin concentration in the lung, there is an increase in total antibody due to the presence of human immunoglobulin in the BALF (**Figure 2 G-L)** from the IgM-IVIg.

### Effect of low dose IgM-IVIg on lung mononuclear phagocytes

Antibodies are capable of directly lysing bacteria however they can also aid clearance by coating the surface of bacteria enhancing uptake by phagocytes in a process known as opsonophagocytosis ^37^. Therefore, we next examined the effects of MCAO and IgM-IVIg treatment on lung mononuclear phagocytes. Flow cytometry was used to detect myeloid cells in the lung (**Figure 5A**). The lung contains two major populations of macrophages, the alveolar macrophage and the interstitial macrophage, although heterogeneity exists within these broad categorisations ^38^. Alveolar macrophages, which are the majority population, were not affected by MCAO or by treatment with IgM-IVIg (**Figure 5B**). In contrast, interstitial macrophages were reduced after MCAO and further reduced after treatment with IgM-IVIg (**Figure 5C**). Interstitial macrophages can be repopulated from blood-derived monocytes ^38^. There was no difference in total monocyte numbers, or Ly6C^lo^ and Ly6C^hi^ functionally distinct monocyte subsets in the lung after MCAO or IgM-IVIg treatment (**Figures 5D, E, F**), suggesting that a lack of precursors was not responsible for the reduction in interstitial macrophages. MCAO also resulted in increased numbers of neutrophils (**Supplementary Figure 4A**) and reduced the number of eosinophils in the lung (**Supplementary Figure 4B**). Lung T cells, dendritic cells and NK cells were unaffected by either MCAO or IgM-IVIg treatment (**Supplementary Figures 4C, F**).

**Figure 5.**
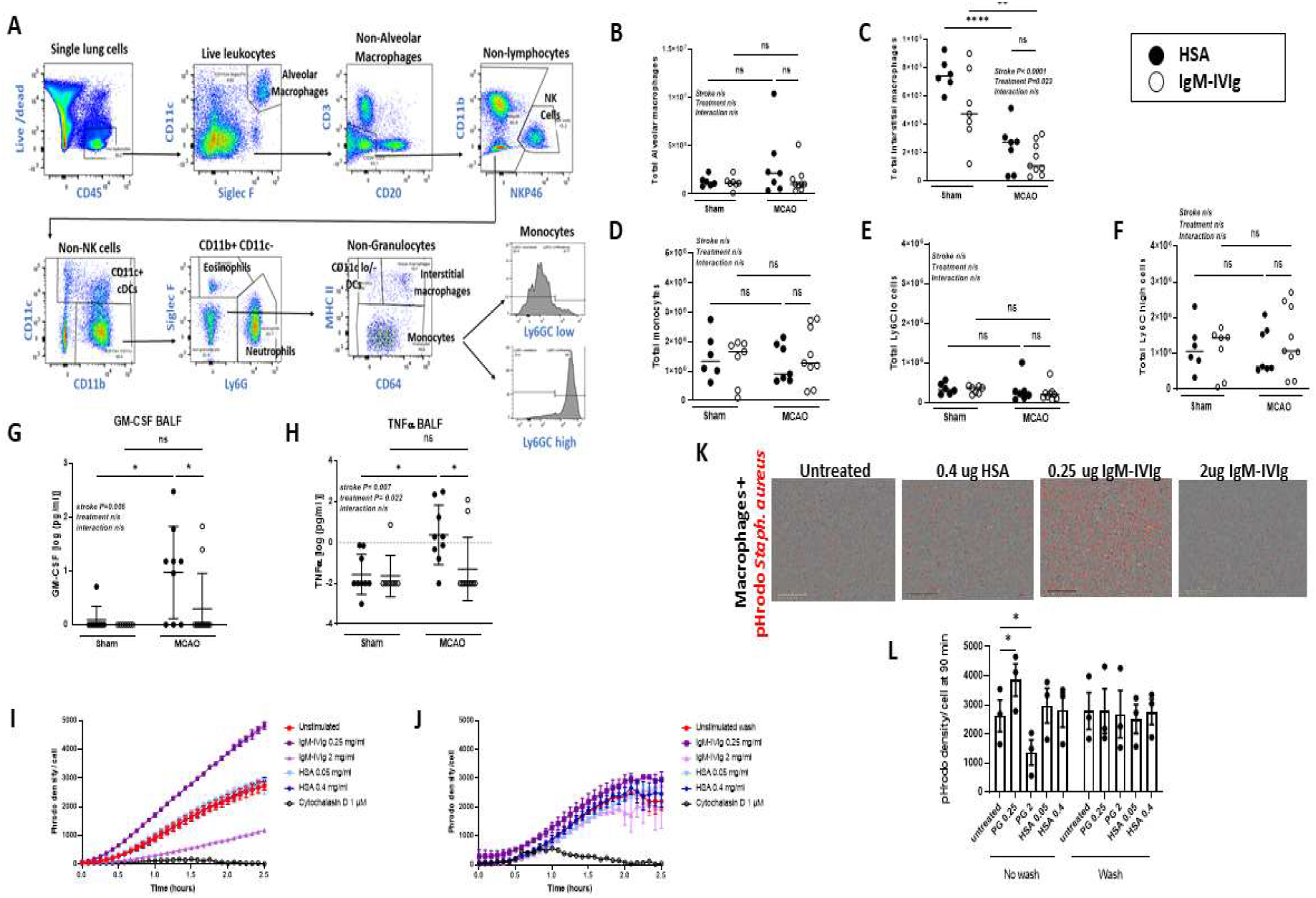
Effect of low dose IgM-IVIg on lung mononuclear phagocytes. (**A**) Gating strategy used to define lung immune cell subsets by flow cytometry. (**B**) Total CD45^+^SiglecF^+^CD11c^+^ alveolar macrophages, (**C**) total CD45^+^CD1b^+^CD64^+^MHCII^+^ interstitial macrophages, (**D**) total CD45^+^CD11b^+^CD64^+^MHC II-monocytes and monocytes distinguished by (**E**) low expression of Ly6C or (**F**) high expression of Ly6C measured by flow cytometry of lung single cell suspensions from mice treated with human serum albumin (HSA; ●) or IgM-IVIg (○) and after 2 d recovery from sham or MCAO surgery (Sham HSA n=6; Sham IgM-IVIg n=7; MCAO HSA n=7; MCAO IgM-IVIg n=10). Concentration of (**G**) GM-CSF (**H**) TNF α measured by multiplexed ELISA of BALF from mice treated with HSA (●) or IgM-IVIg (○) and after 2 d recovery from sham or MCAO surgery (Sham HSA n=9; Sham IgM-IVIg n=8; MCAO HSA n=8; MCAO IgM-IVIg n=11). Phagocytosis of pHrodo® Red S. aureus BioParticles® (fluorescent density/ cell) measured over 2.5 h in co-culture with bone marrow-derived macrophages (BMDM) stimulated for 4 h with IgM-IVIg (0.25 mg/ml and 2 mg/ml) or HSA (0.05 mg/ml and 0.4 mg/ml) where treatments are (**I**) present during phagocytosis assay or (**J**) washed off prior to the addition of pHrodo® Red S. aureus BioParticles®, with cytochalasin D (10μM) as a negative control. Data show technical replicates from one experiment. (**K**) Representative image of phagocytosis of pHrodo® Red S. aureus BioParticles® by BMDM stimulated for 4 h with IgM-IVIg (0.25 mg/ml and 2 mg/ml), HSA (0.4 mg/ml) or cytochalasin D (10 mM) where treatments are present during phagocytosis assay or washed off prior to the addition of pHrodo® Red S. aureus BioParticles®. (**L**) pHrodo® Red S. aureus BioParticles® density/ cell measured after 90 min incubation with bone marrow-derived macrophages stimulated for 4 h with IgM-IVIg (0.25 mg/ml and 2 mg/ml) or HSA (0.05 mg/ml and 0.4 mg/ml) where treatments are present during phagocytosis assay or washed off prior to the addition of pHrodo® Red S. aureus BioParticles®. Data show data points with mean ± S.D; * P<0.05; ** P<0.01; *** P<0.001; P<0.0001; (B-H, L) two way ANOVA with Tukey’s multiple comparison test; (**K**) Scale bar 400 μM.

We next used multiplex ELISA to profile the cytokine environment in the lung after MCAO and treatment with IgM-IVIg. MCAO induced an increase in GM-CSF (**Figure 5G**) and TNF α (**Figure 5H**) concentrations in the BALF, likely a response to post-stroke bacterial pneumonia. Treatment with IgM-IVIg reduced the concentrations of these inflammatory cytokines. Stroke also increased the concentration of IL-33 in the BALF, however this was unaffected by IgM-IVIg treatment (**Supplementary Figure 5A**). There were no significant differences in concentrations of IL-6, IFN γ, IL-1α, Il-2, IL-4, IL-10 and IL-12. However IL-6 and IFN γ showed the same trend of a MCAO-induced increase which was blunted after treatment with IgM-IVIg (**Supplementary Figures 5B-H**).

We next wanted to understand if IgM-IVIg might influence macrophage function, notably, anti-bacterial phagocytic activity given that lung macrophages will be exposed to elevated exogenous immunoglobulin levels after IgM-IVIg treatment (**Figure 2**). Macrophages express multiple Fc receptors which can have both activating and inhibitory effects ^39^, therefore it is plausible that IgM-IVIg could exert immunomodulatory effects on the macrophages themselves which alters their phagocytic capabilities. Furthermore, the presence of intravenous immunoglobulins at high doses can inhibit phagocytosis via saturating macrophage Fc receptors ^40^. To distinguish direct immunomodulatory effects of IgM-IVIg on macrophage phagocytosis from the effects on opsonisation induced uptake of bacteria, we incubated bone marrow-derived macrophages (BMDM) with IgM-IVIg for 4 h. IgM-IVIg was then washed from cells and replaced with media, or left in the well with cells, and pHrodo® Red S. aureus BioParticles® were added. Phagocytosis of pHrodo bioparticles was then measured over 2.5 h. A low dose of IgM-IVIg (0.25 mg/ml) enhanced phagocytosis of pHrodo bioparticles in comparison to control cells that were pre-treated with media only, whereas a high dose of IgM-IVIg (2 mg/ ml) inhibited phagocytosis (**Figure 5G**). Low (0.05 mg/ml) or high (0.25 mg/ml) dose of HSA did not alter phagocytosis above baseline.

In contrast, when IgM-IVIg was washed from cells prior to the addition of pHrodo bioparticles, no effect of IgM-IVIg at low or high dose were observed (**Figure 5H**). The density of internalised pHrodo bioparticles per cell was calculated at 90 min after the addition of pHrodo across three independent experiments. This confirmed low dose IgM-IVIg significantly increased phagocytosis of pHrodo bioparticles when present throughout the assay, however this effect was lost when IgM-IVIg was washed prior to incubation with bioparticles (**Figure 5I, J**).

In conclusion, these data show that reduced bacterial load in the lungs of IgM-IVIg treated mice is not associated with an increased abundance of mononuclear phagocytes within the lung. Certain pro-inflammatory cytokines elevated after MCAO were dampened after treatment with IgM-IVIg. Furthermore, IgM-IVIg does not directly alter the capabilities of macrophages to phagocytose bacteria *in vitro*, but instead at low dose (but not high dose), enhances phagocytosis via bacterial opsonisation.

## DISCUSSION

We describe here the effects of low dose of IVIg (i.e. antibody replacement dose), enriched for IgM in the context of stroke-associated infection. Our key *in vivo* findings are that IgM-IVIg enhances the clearance of bacterial infection and reduces lung pathology and inflammatory cytokines levels alongside systemic B cell composition and antibody levels following experimental stroke. We also observed that IgM-IVIg enhanced bacterial opsonophagocytosis by macrophages *in vitro*, a potential mechanism contributing to enhanced bacterial clearance *in vivo*. Infections are a frequent complication of stroke, affecting up to 30% of stroke patients, and independently affect functional outcome and survival when experienced during the acute and chronic phase of stroke recovery ^5–8^. Bacterial pneumonias have the greatest impact on clinical outcome and are associated also with further complications such as recurrent stroke ^41,42^. Currently there are no available treatments to prevent pneumonia as prophylactic antibiotic treatment after stroke had no effect on pneumonia incidence or clinical outcome in randomised trials ^43,44^. In recent years we have learned that ischemic stroke induces early changes to systemic immune function including atrophy of secondary lymphoid organs, reduced numbers of lymphocytes ^10,12,45–48^ and functional deficiencies in B cell ^12^, T cell ^45^, NKT cell ^49^ and monocyte ^50–52^ populations. Preventing some of the immunosuppressive effects of stroke, or selectively restoring immune function may provide alternative or adjunct therapies to reduce infection incidence and improve outcome and long-term survival after stroke. However, infiltration of certain immune cell populations into the brain and associated inflammatory responses are known to exacerbate brain injury ^53^. Therefore any attempts to enhance immune function to prevent infection must be targeted to prevent unfavourable effects on the developing stroke lesion and processes influencing repair and recovery.

There are 5 main classes of immunoglobulins; IgG, IgM, IgA, IgD and IgE, which all have different properties and functions ^54^. In this study we have used an IgM-enriched preparation of IVIg, Pentaglobin™, which contains 72% IgG, 12%IgM and 16% IgA. This is similar to the proportion of antibodies naturally occurring in serum ^54^, unlike conventional IVIG which is comprised of >96% IgG antibodies. IgM is known to be the most potent antibacterial immunoglobulin due to its highly effective agglutination and complement fixation properties ^55^. Immunosuppressed patients with selective IgM deficiency often present with recurring respiratory infections ^56^. Those with agammaglobulinemia, where production of all immunoglobulin isotypes is impaired, who were given IgG only immunoglobulin replacement therapy had recurrent bacterial lung infections that resulted in chronic lung disease in some ^57^. *In vitro* assays showed Pentaglobin™ had a greater opsonic activity against *Pseudomonas aeruginosa, Staphylococcus aureus* and *Escherichia coli*, in comparison to standard IVIg ^22,27^ which are some of the common bacterial species that cause pneumonia in stroke patients ^58^. Our data, confirm and expand on these findings, demonstrating that at low dose this effect is solely due to opsonisation of bacteria and does not directly influence macrophage phagocytic capacity. However at high dose, the presence of IgM-IVIg actually inhibits this process. In addition to its use in infection prevention, conventional IVIg has also been used at high doses in experimental stroke, where it had immunomodulatory and anti-inflammatory properties and reduced infarct size via supressing TLR responses to damage-associated molecular patterns (DAMPs) and suppressing inflammasome-mediated and complement-mediated neuronal cell death ^59–64^. Therefore it is unlikely that an IVIg immunomodulatory approach after stroke will worsen injury in the brain. In this study, stroke infarct size was unaffected by treatment with low dose IgM-IVIg (**Supplementary Figure 1**), which supported that effects on bacterial load are a direct consequence of its anti-bacterial properties and not due to altering stroke severity. However there may be a dose which confers both anti-infective and brain reparative properties and further studies are warranted to investigate this potential.

Potential harmful effects of IgM-IVIg treatment in the context of cerebrovascular disease also require consideration. IVIg is known to increase serum viscosity and one of the reported adverse effects in a minority population is thromboembolic events, including stroke ^65^. Often IVIg is given as a continuous infusion to minimise effects on blood viscosity. In this study, IgM-IVIg was given as a single bolus pre-surgery and again at 24 h. Treatment had no adverse effects on animal welfare except for a small increase in body weight loss which normalised by 5 d (**Supplementary Figure 1**). Mortality rates were similar between groups at 2 d and improved in IgM-IVIg treated animals at 5 d post experimental stroke (**Supplementary Figure 1**). Our data, together with previous studies, further substantiate the favourable preclinical safety profile of IVIg treatment in stroke.

Low dose IVIg is used as an anti-infective agent in immunosuppressed patients rather than for its immunomodulatory properties reported at high doses. However even at these low doses we saw immunomodulatory effects on plasma B cells and antibodies in the spleen (**Figure 3**). This is in agreement with reports that show IVIg can induce proliferation and immunoglobulin production in B cells from immunosuppressed patients ^66^. Furthermore, IgM-IVIg has β-propiolactone added which reduces complement fixation and prevents aggregation in the circulation reducing systemic adverse effects such as immune complex deposition ^67^. β-propriolactone also reduces binding of immunoglobulins to monocyte Fc receptors reducing the immunomodulatory effects on mononuclear phagocytes ^68^.

Although we have demonstrated important B cell immunomodulatory effects even at this low dose of IgM-IVIg, the main benefits of treatment may be exerted by increasing systemic and lung antibody concentrations, via the provision of human immunoglobulins and the rescue of systemic mouse immunoglobulin concentrations, which in turn improves pathogen opsonisation and clearance. In the lung there was no effect of treatment with IgM-IVIg on plasma B cell number or antibody secretion. It is possible that this is due to the reduced access of exogenous human IgM to the lung compartment versus concentrations that are present in the circulation (**Figure 2**) as signalling via the Fcμ receptor on B cells is known to confer signals for survival and activation ^35^. Furthermore, IgM-IVIg rescued the MCAO-induced loss of the plasma B cell pool in the spleen. There may be no observed effects of treatment as there was no comparable drop in lung plasma B cells after experimental stroke. Indeed lung plasma B cells increased after MCAO, likely in response to the bacterial infection. The increase in plasma B cells observed in the lung after MCAO was not associated with an increase in endogenous mouse immunoglobulins demonstrating that although B cell populations persist in the lung, they may be functionally impaired. Despite this, total lung antibodies were increased by the provision of IgM-IVIg (**Figure 2**) and we demonstrated *ex vivo* that these have the capacity to improve phagocytosis via opsonisation of bacteria. In addition, the mice treated with IgM-IVIg showed lower pathology scores.

This may be in part due to the reduced bacterial loads present in these animals (**Figure 1A-D**), however bacterial load was not significantly associated with pathology score in IgM-IVIg treated animals in contrast to the association observed in controls (**Figure 1 H, I**). This may be in part due to the reduced presence of pro-inflammatory cytokines such as GM-CSF and TNFα which are known to be associated with exacerbated pathology in other respiratory lung diseases, such as SARS-CoV2, and have therefore been proposed as targets for clinical intervention ^69,70^

In conclusion, we propose that treatment with IgM-IVIg offers potential as a safe and effective method to reduce bacterial load and pathology in the lung after stroke. Further preclinical studies are warranted to optimise dosing regimen, including in combination with antibiotics, and at delayed time points after MCAO to better understand the translational potential of IgM-IVIg as an anti-infective agent for stroke patients. Modulation of dose may be able to harness direct systemic anti-bacterial and neuroimmunomodulatory properties to result in a treatment with dual benefits by preventing infection and reducing harmful aspects of brain inflammation in stroke patients.

## Acknowledgements

We thank staff in the University of Edinburgh biological research facility on the Edinburgh Bioquarter campus for animal husbandry and technical support, in particular Will Mungall who performed all of the i.v. tail vein injections of treatment. We thank staff in the QMRI Flow Cytometry and cell sorting facility (University of Edinburgh) for their support in generating the flow cytometry data and staff at Shared University Research facilities (SURF; University of Edinburgh) for their help with processing, embedding and assistance with slide scanning and Easter Bush Pathology Laboratory (University of Edinburgh) for cutting and H&E staining of lung sections. This work was funded by a grant from the Medical Research Council (Grant Number MR/R001316/1). L.M. is supported by a Sir Henry Dale Fellowship jointly funded by the Wellcome Trust and the Royal Society (Grant Number 220755/Z/20/Z). J.R.G is funded by a Senior Fellowship awarded by The Kennedy Trust for Rheumatology Research. B.W.M., S.M.A. and C. J. S. receive funding from Leducq Foundation

Transatlantic Network of Excellence, Stroke-IMPaCT (Grant Number 19CVD01), and B.W.M receives funding from the UK Dementia Research Institute which receives its funding from the Medical Research Council, Alzheimer’s Society, and Alzheimer’s Research UK.

## Author contributions

Conceptualization, L.M., S.M.A., C.J.S., and B.W.M..; methodology, L.M., A.J.H., A.M., M.J.D.D., S.M.A., C.J.S. and B.W.M.; validation, L.M., A.J.H., A.M., M.J.D.D., M.Y., J.R.G., S.M.A., C.J.S. and B.W.M.; formal analysis, L.M., A.J.H., A.M., and B.W.M.; investigation, L.M., A.J.H., A.M., and M.J.D.D.; resources, L.M., J.R.G., S.M.A., C.J.S., and B.W.M.; data curation, L.M., A.J.H., A.M., and B.W.M. writing – original draft, L.M. and B.W.M..; writing – review & editing, L.M., A.J.H., A.M., M.J.D.D., M.Y., J.R.G., S.M.A., C.J.S. and B.W.M. ; visualization, L.M., A.J.H., A.M., and B.W.M. ; supervision, L.M., S.M.A., C.J.S., and B.W.M; project administration, L.M., J.R.G., S.M.A., C.J.S., and B.W.M.; funding acquisition, L.M., J.R.G., S.M.A., C.J.S and B.W.M.

## Declaration of Interest

No competing interests to declare.

---

**Supplementary Figure 1.**
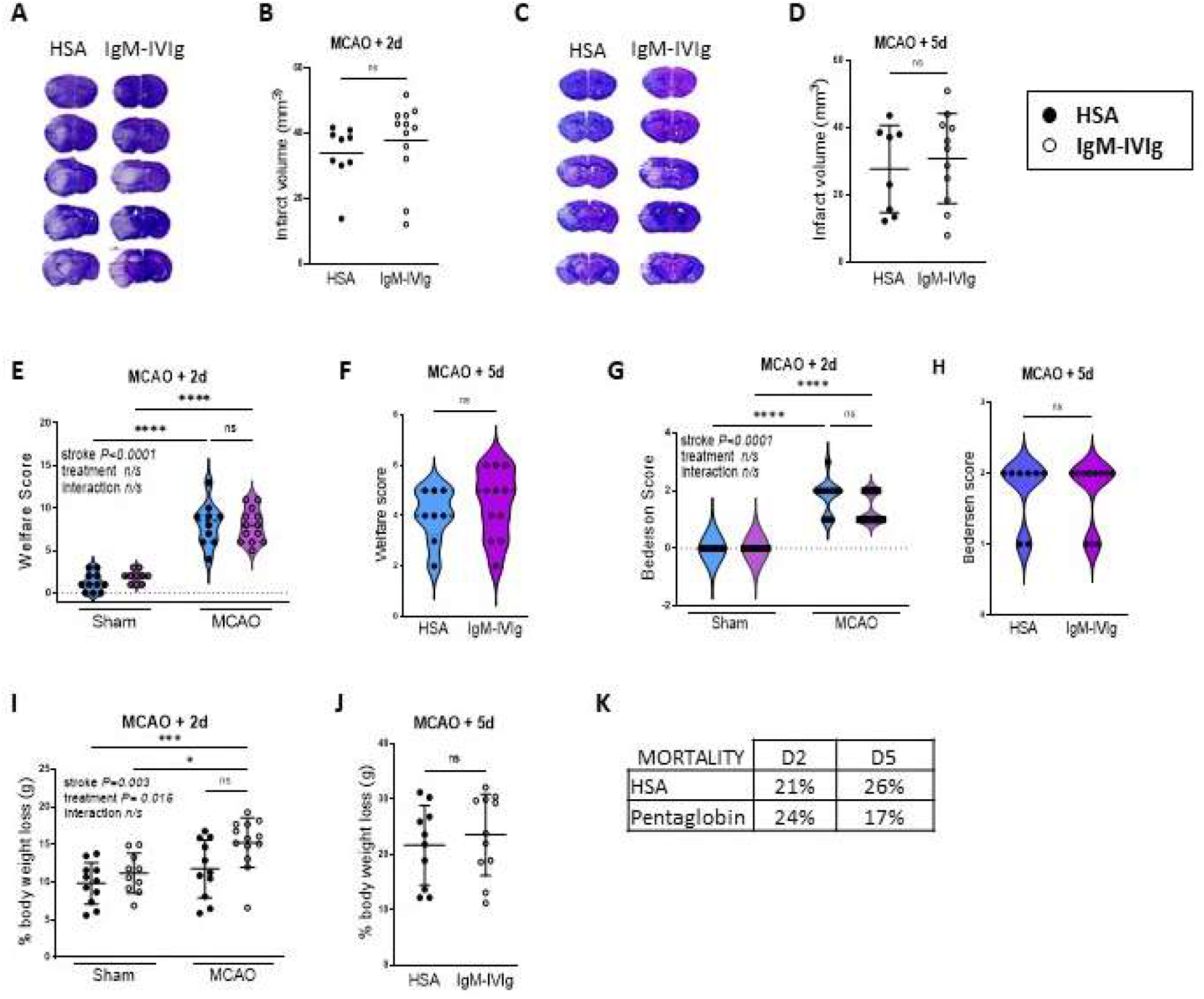
Low dose IgM-IVIg does not modulate primary stroke pathology. Infarct volume (mm^3^) in the brains of animals treated with human serum albumin (HSA; ●) or IgM-IVIg (○) at (**A, B**) 2 days or (**C,D**) 5 days post MCAO. Treatment with IgM-IVIg did not affect welfare score at (**E**) 2 days or (**F**) 5 days after MCAO. (**G**) Animals treated with IgM-IVIg had a higher percentage reduction in bodyweight 2 days post sham or MCAO surgery. (**H**) However at 5 days post MCAO there was no effect of treatment with IgM-IVIg on percentage body weight loss. Treatment with IgM-IVIg did not affect Bedersen score at (**K**) 2 days or (**I**) 5 days after MCAO. (**J**) Treatment with IgM-IVIg had no effect on mortality at 2 days after MCAO however at 5 days post MCAO, mortality was lower in mice treated with IgM-IVIG (**B, D, G, H, K, L**) Data show data points with mean ± S.D; * P<0.05; ** P<0.01; *** P<0.001; P<0.0001; (**B, D, H, L**) unpaired t-test. (**G, K**) two way ANOVA with Tukey’s multiple comparison test; (**E, F, I, J**) Violin plots with Mann-Whitney

**Supplementary Figure 2.**
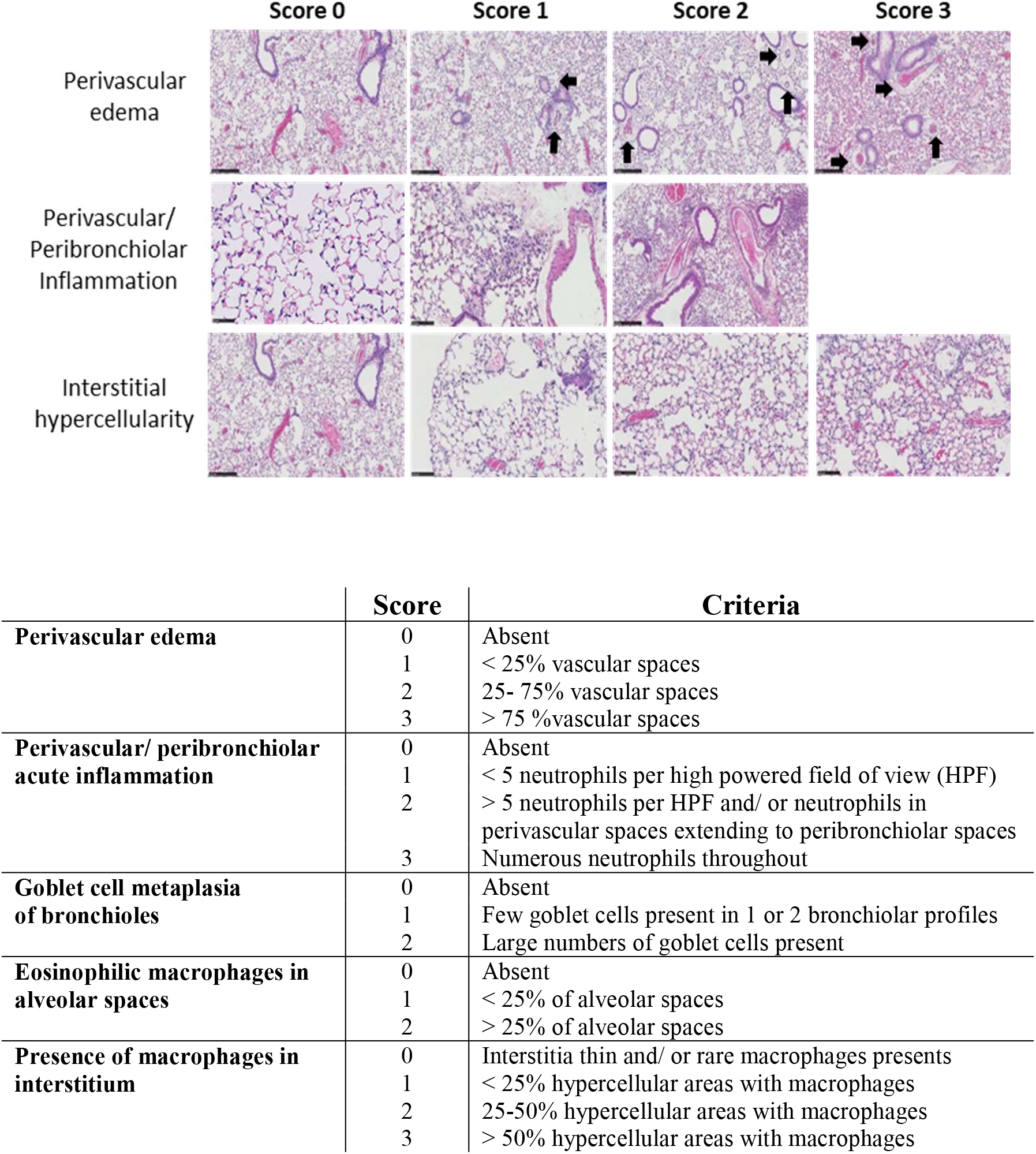
Criteria for scoring lung pathology. Scoring system used to grade the extent of pathology in lung from animals treated with HSA or IgM-IVIG 2 days and 5 days after sham or MCAO surgery (**Figure 1**). Goblet cell hyperplasia was not detected in any animals. All animals had <25% alveolar spaces with macrophages present and scored 1. Inflammation did not reach a score of 3 in any animals. Scores in sham-operated animals mainly consisted of edema and some with mild inflammation. Scale bars 50 mm.

**Supplementary Figure 3.**
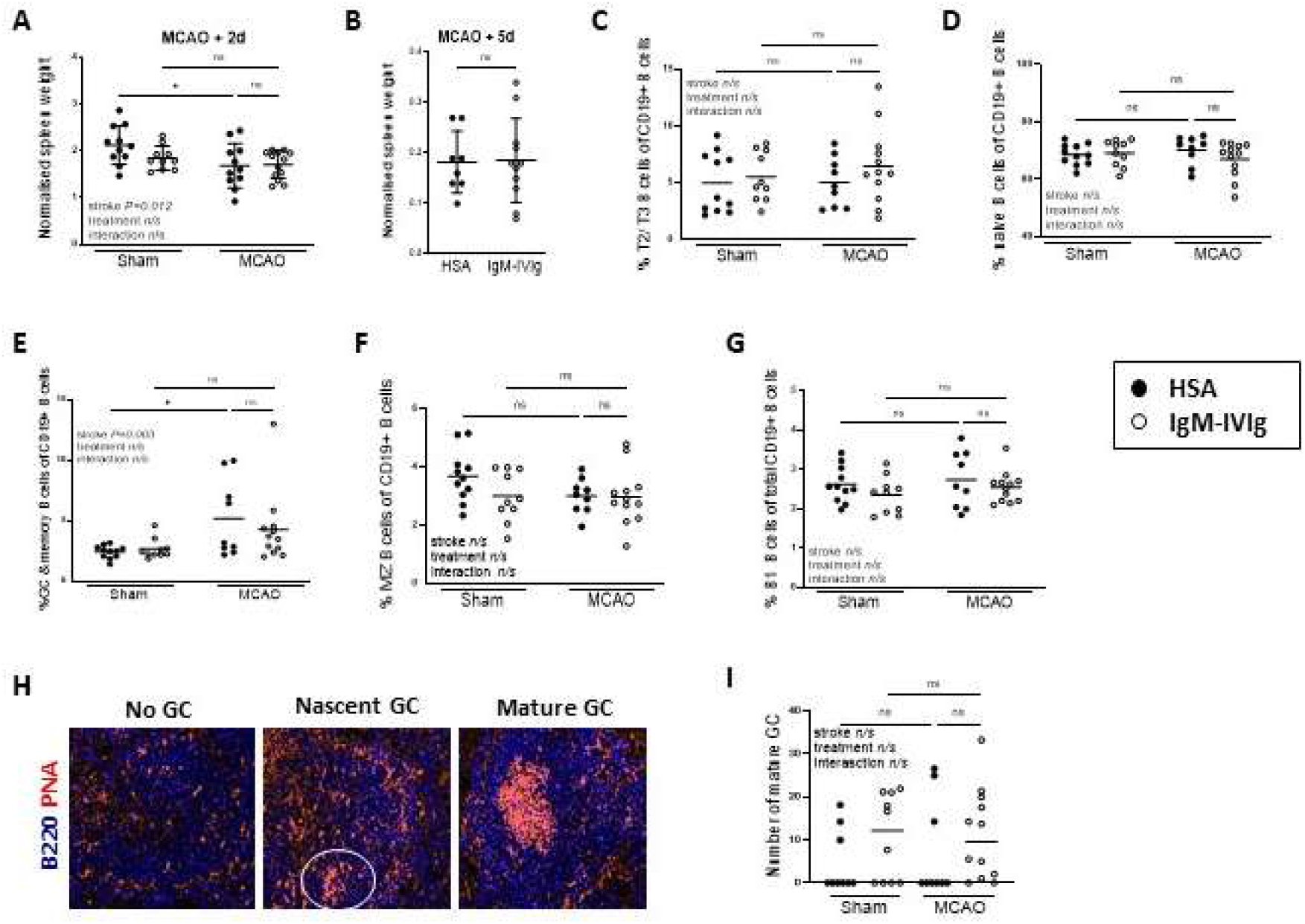
Splenic B cell responses to IgM-IVIg. Spleen weight, normalised to total body weight, is significantly reduced by stroke (**A**) but there is no effect of treatment with IgM-IVIg at (**A**) 2 d or (**B**) 5 d after sham or MCAO surgery. Percentage of (**C**) IgD^+^CD23^+^CD24^+^CD93^+^ T2 transitional B cells (**D**) IgD^+^CD23^+^CD23^+^CD24^+^CD93^−^ naïve B cells (**E**) CD23^−^CD21^−^CD93^−^ GC and memory B cells (**F**) CD19^+^CD23^−^CD21^+^CD93^−^ marginal zone B cells and (**G**) CD93^−^CD43^+^ B1 B cells within the total CD19^+^ B cell population measured by flow cytometry of spleens from mice treated with human serum albumin (HSA; ●) or IgM-IVIg (○) and after 2 d recovery from sham or MCAO surgery. Gating strategy in **Figure 3A** (Sham HSA n=11; Sham IgM-IVIg n=10; MCAO HSA n=9; MCAO IgM-IVIg n=12). (**H**) Immunolabelling of germinal centres (GC) using peanut agglutinin (PNA; orange) and DAPI staining of nuclei (blue) in spleens to identify areas of white pulp with no GC, nascent GC and mature GC. (**I**) Number of mature GC per half spleen section in spleens from mice treated with human serum albumin (HSA; ●) or IgM-IVIg (○) and after 2 d recovery from sham or MCAO surgery (Sham HSA n=9; Sham IgM-IVIg n=10; MCAO HSA n=9; MCAO IgM-IVIg n=12). Data show data points with mean ± SD; (**A, B, C, D, E , G**) two way ANOVA with Tukey’s multiple comparison test

**Supplementary Figure 4.**
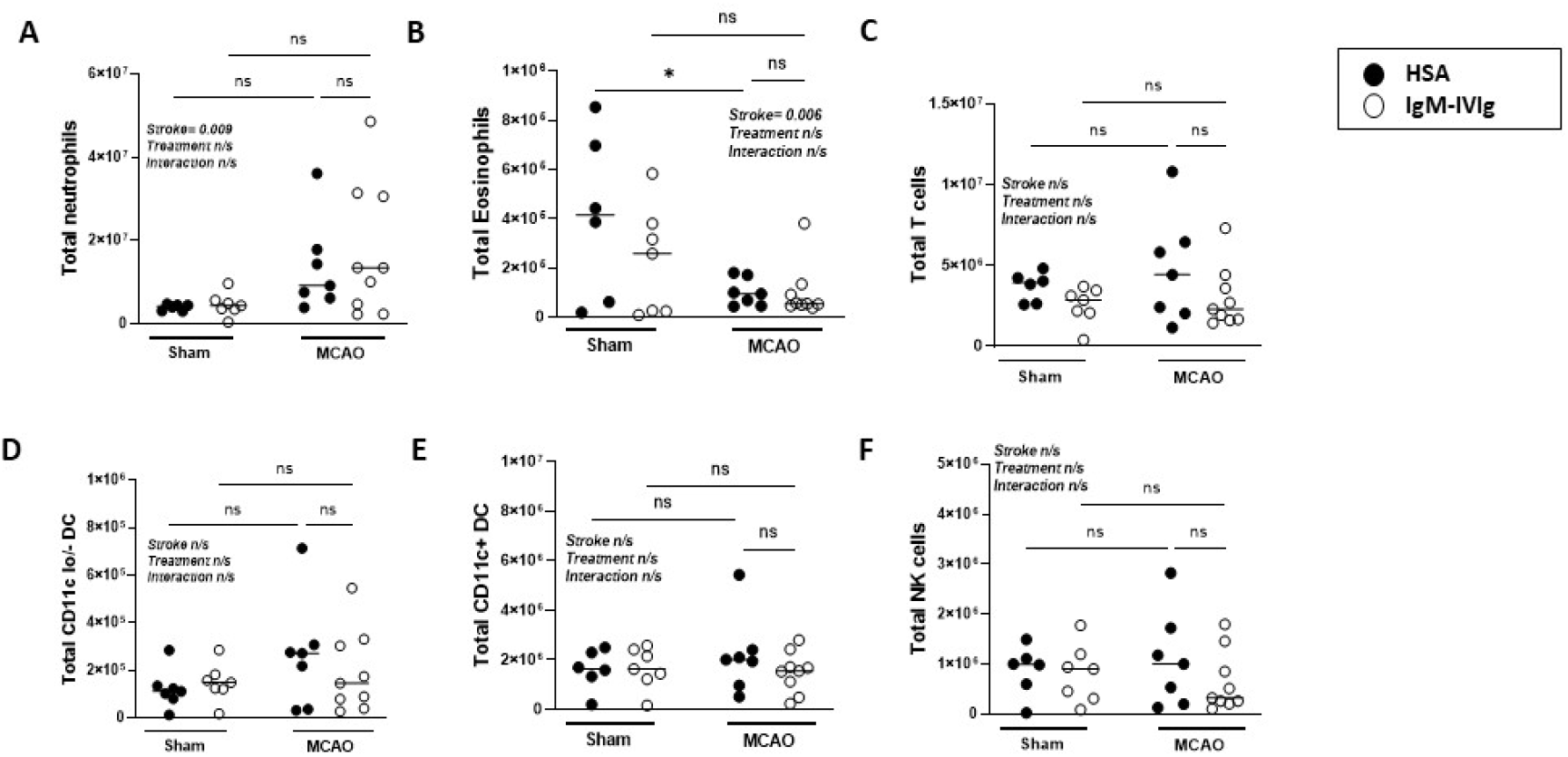
Effect of IgM-IVIg on lung immune cell subsets. Total (**A**) CD11b+SiglecF-LygG^+^ neutrophils, (**B**) CD11b^+^SiglecF^+^LygG^−^ eosinophils, (**C**) CD3^+^CD20^−^ T cells, (**D**) CD11b^+^SiglecF^−^Ly6G^−^CD64^−^MHC II^+^ DC (**E**) CD11b ^+/-^CD11c^+^ DC and (**F**) CD11b^−^NKP46^+^ NK cells measured by flow cytometry of lung single cell suspensions from mice treated with human serum albumin (HSA; ●) or IgM-IVIg (○) and after 2 d recovery from sham or MCAO surgery. For gating strategy see **Figure 4A**. (Sham HSA n=6; Sham IgM-IVIg n=7; MCAO HSA n=7; MCAO IgM-IVIg n=10). Data show data points with mean ± S.D; * P<0.05; (**A-F**) two way ANOVA with Tukey’s multiple comparison test.

**Supplementary Figure 5.**
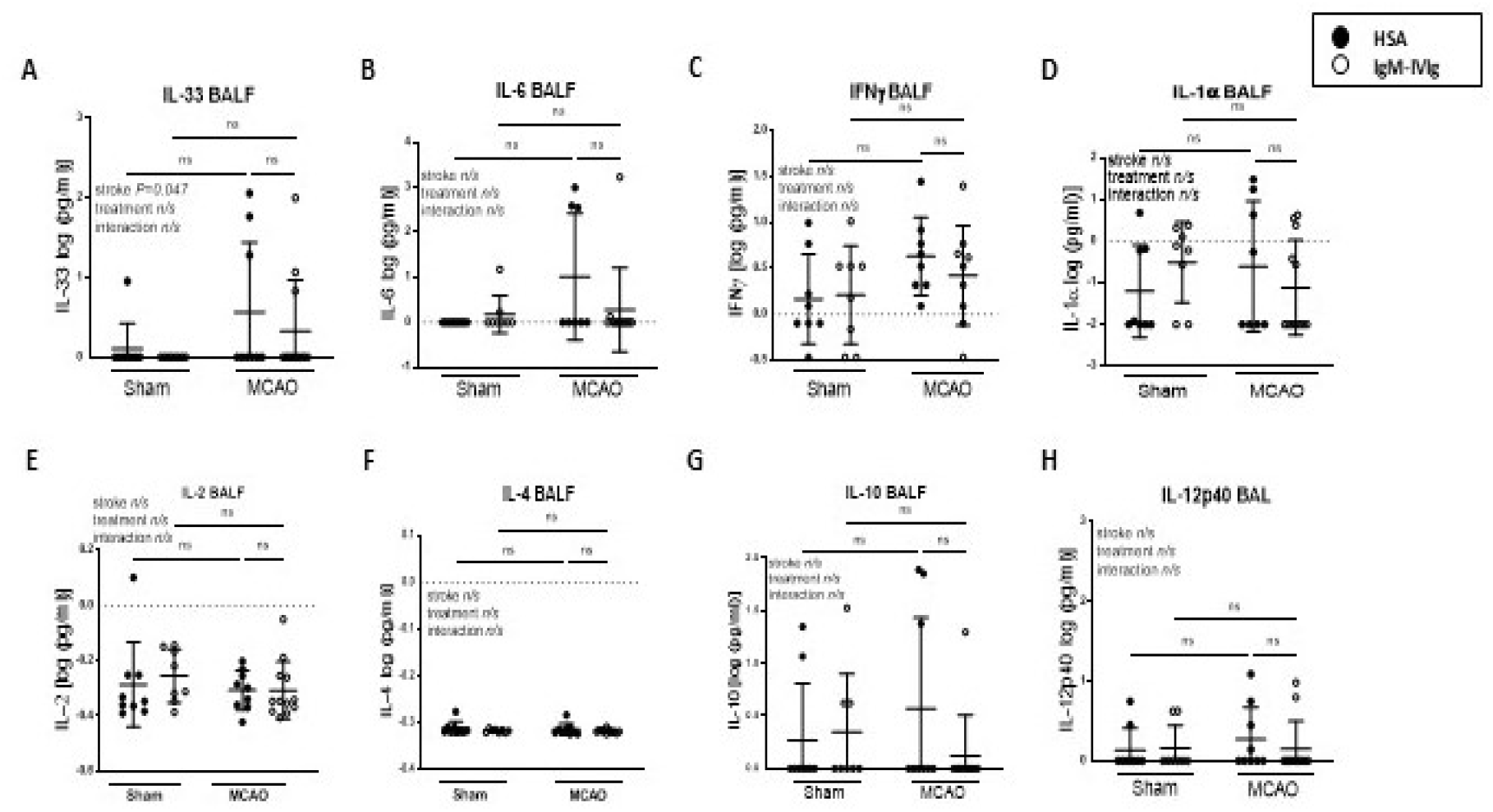
Effect of IgM-IVIg on lung immune cell subsets. Concentration of (**A**) IL-1a (**B**) IL-2 (**C**) IL-4 (**D**) IL-6 (**E**) IL-10 (**F**) IL-12p40 (**G**) IL-33 and (**H**) IFNg measured by multiplexed ELISA of BALF from mice treated with human serum albumin (HSA; ●) or IgM-IVIg (○) and after 2 d recovery from sham or MCAO surgery (Sham HSA n=9; Sham IgM-IVIg n=8; MCAO HSA n=8; MCAO IgM-IVIg n=11). Data show data points with mean ± S.D; (**A-H**) two way ANOVA with Tukey’s multiple comparison test.

## Notes

### Competing Interest Statement

The authors have declared no competing interest.

